# Hybrid incompatibility between *D. virilis* and *D. lumei* is stronger in the presence of transposable elements

**DOI:** 10.1101/753814

**Authors:** Dean M. Castillo, Leonie C. Moyle

## Abstract

Mismatches between parental genomes in selfish elements are frequently hypothesized to underlie hybrid dysfunction and drive speciation. However, because the genetic basis of most hybrid incompatibilities is unknown, testing the contribution of selfish elements to reproductive isolation is difficult. Here we evaluated the role of transposable elements (TEs) in hybrid incompatibilities between *Drosophila virilis* and *D. lummei* by experimentally comparing hybrid incompatibility in a cross where active TEs are present in *D. virilis* (TE+) and absent in *D. lummei*, to a cross where these TEs are absent from both *D. virilis* (TE-) and *D. lummei* genotypes. Using genomic data, we confirmed copy number differences in TEs between the *D. virilis* (TE+) strain and both the *D. virilis* (TE-) strain and *D. lummei*. We observed F1 postzygotic reproductive isolation exclusively in the interspecific cross involving TE+ *D. virilis* but not in crosses involving TE- *D. virilis*. This mirrors intraspecies dysgenesis where atrophied testes only occur when TE+ *D. virilis* is the paternal parent. A series of backcross experiments, that accounted for alternative models of hybrid incompatibility, showed that both F1 hybrid incompatibility and intrastrain dysgenesis are consistent with the action of TEs rather than genic interactions. Thus, our data suggest that this TE mechanism manifests as two different incompatibility phenotypes. A further Y-autosome interaction contributes to additional, sex-specific, inviability in one direction of this cross combination. These experiments demonstrate that TEs that cause intraspecies dysgenesis can increase reproductive isolation between closely related lineages, thereby adding to the processes that consolidate speciation.

## Introduction

Selfish elements have been proposed to play a key role in the evolution of reproductive barriers between species. However, most evidence for this role in both plants and animals comes from empirical studies that indirectly correlate differences in repetitive elements between species with both increased transposable element (TE) transcription and phenotypic dysfunction in hybrids (Labrador *et al*. 1999; Martienssen 2010; Brown *et al*. 2012; Dion-Cote *et al*. 2014). In comparison, there have been few attempts to more directly assess the role of TEs in the manifestation of reproductive isolation, using patterns of hybrid dysfunction in controlled crosses, especially in comparison to genic differences between lineages. Early attempts to do so concluded that TEs did not contribute to reproductive isolation because dysfunctional phenotypes typically associated with TE mobilization, and increased mutation rates (a signature of TE movement), were not observed in several species crosses that otherwise have marked postzygotic isolation (Coyne 1986; Hey 1988; Coyne 1989). Differences between these species in TE copy number for specific elements and TE families/subfamilies were not estimated. This inferred lack of a strong direct effect of TEs on between species incompatibility contrasted with strong evidence in some species groups for TEs contributing to intraspecies dysgenesis: an asymmetrical incompatibility among strains where the paternal genotype carries copies of a specific TE element whereas the maternal genotype lacks copies of this element (Kidwell 1985). We now understand that these intraspecific dysgenesis phenotypes reflect more complex effects of de-repression of TE transcription (Martienssen 2010; Khurana *et al*. 2011) and genome wide DNA damage (Khurana *et al*. 2011). This mismatch between parental genomes is mechanistically connected to these dysgenesis phenotypes through the machinery that regulates the silencing of TEs in the genome. In most eukaryotic genomes, TEs are epigenetically silenced through a small-RNA-mediated process that heterochromatizes these loci. These small-RNAs are maternally loaded into the embryo and are necessary for TE silencing (reviewed in Slotkin and Martienssen 2007). When small-RNAs are not loaded into the embryo by a maternal parent because they lack copies of a TE, the resulting zygote is susceptible to the negative consequences of TE misregulation.

This potential to cause negative genetic interactions via TE de-repression in hybrids clearly places TEs within the classical Dobzhansky-Muller framework for the evolution of incompatibilities (Dobzhansky 1937; Muller 1942)—whereby postzygotic isolation results from negative epistasis among divergent loci in hybrid genomes (Johnson 2010; Maheshwari and Barbash 2011; Castillo and Moyle 2012; Crespi and Nosil 2013). Moreover, we now also have the means to better assess differences in TE identity, copy number, and age, between closely-related lineages and species via high throughput DNA and RNA sequencing. These data, and new analytical tools to estimate copy number, make it timely to revisit the relative contribution of TEs to reproductive isolation phenotypes using dissection with experimental crosses, just as has been shown with standard genic incompatibilities (Johnson 2010; Presgraves 2010; Castillo and Barbash 2017).

The observation that TE content frequently differs between closely-related lineages, both between species and among strains within a species (Kidwell and Lisch 1997; Stuart et al. 2016; Petersen et al. 2019) also provides an experimental mechanism for addressing the contribution of TEs to species reproductive barriers. In particular, when strains within species differ in the presence of active TEs, inter-specific crosses involving these strains are predicted *a priori* to differ in their magnitude of hybrid dysfunction. Specifically, stronger barriers are expected in crosses involving strains containing active TEs compared to those without active TEs. Of known instances of intraspecific TE polymorphism, dysgenic systems in *Drosophila* are also among the best characterized in terms of the deleterious phenotypic effects of TEs in between-strain crosses, that could also be relevant to reproductive isolation between species. In particular, the dysgenic syndrome within *Drosophila virilis* and the *P-element* system within *D. melanogaster* both typically exhibit gonadal atrophy in offspring when a strain carrying TEs is crossed with a strain lacking these elements (Kidwell 1985; Lozovskaya *et al*. 1990). The dysgenic phenomenon depends both on the direction of the cross—dysgenesis occurs when the female parent lacks the TE elements—and on the copy number in the paternal parent (Srivastav and Kelleher 2017; Serrato-Capuchina *et al*. 2020b). These intraspecific dysgenic systems can be used to evaluate the contribution of TEs to reproductive isolation with other species, when there are two strains that differ in the presence of TEs known to cause intraspecific dysgenesis and a second closely related species that lacks these TEs. If the presence of TEs amplifies hybrid incompatibility between species, an interspecific cross involving the TE+ carrying strain should exhibit greater hybrid incompatibility than the same interspecific cross but using a strain in which TEs are absent.

In this study, we use this logic to evaluate whether TEs affect the magnitude of reproductive isolation between *D. lummei* and *D. virilis*—the latter of which is well known for its intraspecific dysgenic system (Lozovskaya *et al*. 1990). These and other species in the virilis clade are closely related and variable in the degree to which they exhibit premating and postmating reproductive isolation. Importantly, they also vary in TE copy number of specific elements (Zelentsova *et al*. 1999; Evgen’ev *et al*. 2000). Dysgenesis within *D. virilis* was first described by Lozovskaya and colleagues (Lozovskaya *et al*. 1990) when they observed gonadal atrophy and sterility in males and females produced in crosses between strains that varied in TEs. As expected, *D. virilis* has large among-strain variation in the copy number of active TE elements (below). In contrast, *D. lummei* has a different profile of active TEs compared to *D. virilis* (Zelentsova *et al*. 1999; Evgen’ev *et al*. 2000; see below), and is not reported to exhibit intraspecific dysgenesis.

Using the features of this system and the established behavior of dysgenesis among *D. virilis* strains, we contrast two crosses where parental strains vary in their differences in dysgenic TE copy number to evaluate the connection between postzygotic reproductive isolation and differences in TE composition. Specifically we compare a cross where dysgenic TEs are present in the *D. virilis* (TE+) parent and absent in the *D. lummei* parent to a cross where dysgenic TEs are absent from both *D. virilis* (TE-) and *D. lummei* parental genotypes (Fig. 1). In intraspecific crosses between female *D. virilis* that lack relevant TEs (TE-) and male *D. virilis* that carry these TEs (TE+), strong dysgenesis is observed. Therefore, in interspecific crosses we expect the strongest reproductive isolation specifically between *D. lummei* females that lack dysgenic inducing TEs and *D. virilis* males that are TE+ (Fig. 1). With these expectations we show, using a series of directed crosses and backcrosses, that patterns of reproductive isolation observed in our study are consistent with a causal role for TEs increasing reproductive isolation between species.

**Figure 1.**
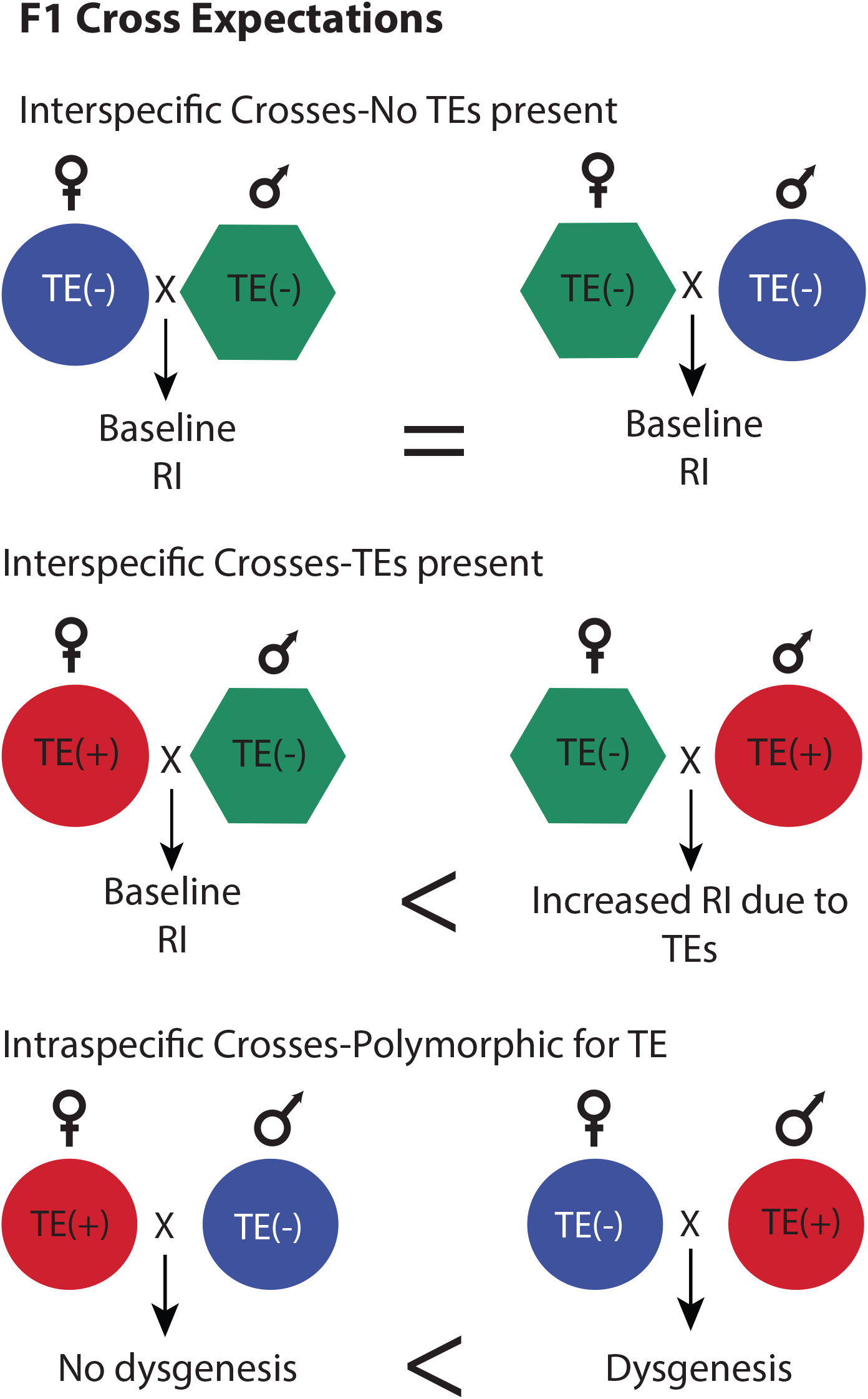
We hypothesize that transposable elements (TEs) will cause increases in reproductive isolation compared to interspecies crosses where both strains lack TEs. The pattern of increased reproduction should be asymmetrical, with the highest level observed when the paternal species carries TE copies that are absent in the maternal species.

## Materials and Methods

### Fly stocks

Our experiments used two *D. virilis* stocks that are known to differ in their copy number of specific TEs, and one stock of the closely related species *D. lummei*. We chose stocks that mated readily to facilitate genetic analysis. The *D. virilis* Strain 9 is a wild type strain that was collected in Georgia, USSR in 1970; it lacks TEs that induce dysgenesis in intraspecific crosses (Lozovskaya *et al*. 1990; Blumenstiel and Hartl 2005). Here we refer to this strain as TE-. The reference genome strain (UCSD stock number 15010-1051.87) contains TEs not present in the TE- strain, and as a result induces dysgenesis in intraspecies crosses (Blumenstiel 2014); we refer to this strain as TE+. The TE- and TE+ strains differ at more than just presence/absence of TEs; for example, one study estimates ~7.35 SNP/kb differentiate them (Hemmer *et al*. 2020). This level of genetic differentiation is consistent with that expected between strains within a diverse species. Accordingly, prior to our experiment we had no expectation that these two strains of *D. virilis* had differentially accumulated other alleles (outside of TE abundance) that also contribute to postzygotic isolation in crosses with *D. lummei*. The reference genome strain of *D. virilis* was used instead of *D. virilis* Strain 160, which is also commonly used in dysgenic studies, because initial crosses between Strain 160 and *D. lummei* showed significant premating isolation in single pair crosses. The reference genome strain and Strain 160 have similarly high levels of TEs (Lozovskaya *et al*. 1990 and results below). Our strain of *D. lummei* was acquired from the UCSD stock center (15010-1011.07).

#### Estimating copy number of dysgenic causing elements

To examine the copy number of potentially dysgenic causing elements, we used publicly available genomes from several *D. virilis* strains and other species in the virilis clade, including two *D. lummei* strains. The SRA accession numbers and strain information for all samples are listed in Supplemental Table 1. We focused on candidate TEs that are associated with dysgenesis (Funikov *et al*. 2018) but mapped reads to all inserts from a previously compiled TE library (Erwin *et al*. 2015) to prevent bias that might arise from mapping reads from the other species to a TE library generated from *D. virilis*. All copy number estimates were made using deviaTE (Weilguny and Kofler 2019), which estimates haploid copy number based on single-copy “control” genes. For our control genes we chose the *D. virilis* homologs of Beta-tubulin60D and alpha-tubulin84B as essential single copy genes in the *Drosophila virilis* genome.

We first compared copy number differences between the genomes of the two strains typically used in studies of dysgenesis: inducing strain *D. virilis* Strain 160 and *D. virilis* Strain 9 (TE-). We determined the reliability of our estimates for identifying potentially causal TEs, by comparing our estimate for the enrichment of TE copy number in Strain 160 to other studies that have used read mapping copy number estimates from genomic data and copy number estimates for piRNA from ovaries of each of these strains (Erwin *et al*. 2015; Funikov *et al*. 2018). Up until recently, the majority of work on dysgenesis in *D. virilis* had focused on a specific retroelement—*Penelope*—as the likely causal element of inter-strain dysgenesis, based on correlations between dysgenic phenotypes and the presence of *Penelope* piRNA (Blumenstiel and Hartl 2005) and *Penelope* copy number (Veiera *et al*. 1998). More recent evidence, however, has excluded a direct causal role for Penelope in dysgenesis (Rozhkov *et al*. 2013; Funikov *et al*. 2018). Nonetheless because we know that dysgenesis is copy number dependent (Veiera *et al*. 1998; Serrato-Cappuchina *et al*. 2020b), we predicted that we could identify potentially causative TEs by determining which other TEs had similar copy number distribution compared to *Penelope*. To do so, we looked at relationships between candidate TEs and *Penelope* across a sample of *D. virilis* genomes. We specifically focused on *Polyphemus, Paris, Helena, Skippy*, and *Slicemaster*, given their greater copy number estimates in inducer vs noninducer strains based on our (below) and previous (Funikov *et al*. 2018) analyses. We used hierarchical clustering to determine which of these candidate TEs had the most similar copy number distribution compared to the *Penelope* element, across all of the *D. virilis* genomes. Any candidate TE that clustered with (showed high similarity to) *Penelope* is inferred to be potential causal of dysgenesis because its copy number variation should be similarly associated with variation in dysgenesis. Specifically, we expected a higher copy number in the TE+ strain compared to the TE- strain and a positive correlation between any causative TE and *Penelope* because a higher copy number would be required to observe asymmetrical dysgenesis in crosses between these strains.

We next determined which candidate elements had a higher average copy number in the two inducing strains (the genome strain TE+ and strain 160) compared to the non-inducing Strain 9 (TE-) and to the copy number average of two *D. lummei* strains for which we also had genome data. We retained the threshold of two-fold increase in the average TE copy number, based on the analysis above. We also examined the differences in the number of polymorphic sites and allele frequencies at each position in the consensus sequence in the *D. lummei* strains compared to *D. virilis* strains, to infer whether TEs were recent/active versus older/inactive. To do so, we filtered SNPs using default settings in deviaTE to calculate the average major allele frequency at each site as well as the number of sites that were polymorphic. TEs that reflect either new invasions or ongoing activity will have a distinct pattern of sequence variation—fewer polymorphic sites, and (at these sites) one or few dominant alleles—resulting in copies across the genome that are highly homogenous. In contrast, the observation of numerous variable sites across element copies indicates older inactive elements (Beall *et al*. 2002; Erwin *et al*. 2015; Weilguny and Kofler 2019).

### Intra- and interspecies crosses to determine the nature of reproductive isolation

To compare intra- and interspecific hybrid phenotypes, we used a design that was parallel for all experimental crosses done within and between species. For all our crosses, virgin males and females from each stock were collected as they eclosed and aged for seven days prior to matings. All crosses involved single pairs of the focal female and male genotype combination (a minimum of 20, range 20-23, replicates were completed for a given female × male genotype combination). We performed three intra-strain crosses (TE- × TE-, TE+ × TE+, and *D. lummei* × *D. lummei*) to account for intrinsic fecundity differences in subsequent analyses of inter-strain crosses. As expected, in the three intra-strain crosses we did not see any evidence of male or female gonad atrophy (classical dysgenic phenotypes) or skewed sex ratios. There were differences in the total number of progeny produced, hatchability, and proportion of embryos that had reached the pre-blastoderm stage (hereafter ‘proportion fertilized’). We used these values as baseline estimates of maternal effects in our statistical models, to account for their potential influence on these phenotypes in our subsequent inter-strain analyses (Table 1; Table 2).

**Table 1.** Average trait means for control intrastrain crosses. These values are used as baseline in statistical models testing for increased reproductive isolation in interspecis crosses. For male dysgensis (n) refers to the number of male progeny scored. se=standard error. The number of replicates used to calculate the standard error are n=22 (TE-), n=23 (TE+), and n=20 (*D. lummei*)

| Female | Male | Dysgenesis (n) | Sex Ratio (se) | Viability (se) |
| --- | --- | --- | --- | --- |
| TE- | TE- | 0.0 (1398) | 0.488 (0.016) | 0.844 (0.055) |
| TE+ | TE+ | 0.0 (575) | 0.516 (0.014) | 0.571 (0.057) |
| *D. lummei* | *D. lummei* | 0.0 (335) | 0.521 (0.015) | 0.411 (0.067) |

**Table 2.** F1 postzygotic reproductive isolation, as observed in crosses between TE+ and *D. lummei*, is a product of embryo inviability. Inviability is inferred when fewer eggs hatch than expected based on the proportion of eggs that initiated development (proportion fertilized). The samples sizes (n) are the pooled number of embryos examined across replicate crosses. † represents a significant difference in the proportion fertilized for an interspecific cross compared to the control cross of the maternal strain (*P*<0.05).

| Female | Male | Proportion Fertilized (n) | Proportion Hatched (n) | $\chi^2_1$ (P-value) |
| --- | --- | --- | --- | --- |
| TE- | TE- | 0.85 (179) | 0.79 (344) | 2.76 (0.09) |
| TE+ | TE+ | 0.58 (155) | 0.55 (224) | 0.21 (0.65) |
| *D. lummei* | *D. lummei* | 0.61 (218) | 0.67 (260) | 1.75 (0.18) |
| TE- | TE+ | 0.85 (180) | 0.80 (283) | 1.00 (0.31) |
| TE+ | TE- | 0.57 (153) | 0.57 (206) | 0.00 (1.00) |
| TE- | *D. lummei* | 0.33† (389) | 0.39 (79) | 0.59 (0.44) |
| *D. lummei* | TE- | 0.60 (161) | 0.75 (337) | 12.18 (<0.001) |
| TE+ | *D. lummei* | 0.55 (126) | 0.42 (235) | 5.41 (0.02) |
| *D. lummei* | TE+ | 0.43† (212) | 0.31 (208) | 6.06 (0.01) |

For inter-strain comparisons, we performed all reciprocal crosses between the three strains to analyze reproductive isolation in F1 progeny. These crosses fall into three categories: **I-Interspecies crosses with no TEs present:** Crosses between TE- and *D. lummei* were used to determine the nature of reproductive isolation between the species, in the absence of dysgenic inducing TEs. **II-Interspecies crosses with TEs present:** Crosses between TE+ and *D. lummei* were used to determine the additional contribution, if any, of TEs to interspecific post-mating reproductive isolation (Fig. 1). This contrast assumes that the two *D. virilis* strains, TE- and TE+, do not differ from each other in their genic Dobzhansky-Muller incompatibilities with *D. lummei*, an assumption that we evaluated with data from subsequent crosses performed using hybrid genotypes (described further below). **III-Intraspecific Crosses Polymorphic for TEs:** Crosses between TE- and TE+ were used to confirm the expression of intraspecific male dysgenesis when the inducing (TE+) strain was used as a father, as well as the level of rescue observed when F1 males were used as male parents crossed to the TE- strain. This cross allowed us to draw parallels between interspecific reproductive isolation and intraspecific dysgenesis.

### Backcross design to test models of hybrid incompatibility

The phenotypes that we observed for postzygotic reproductive isolation could be the product of several non-mutually exclusive genetic mechanisms, including both TEs and genic interactions. Therefore we used a set of backcross experiments to compare the fit of several alternative models of hybrid incompatibility to our data for interspecific reproductive isolation and *D. virilis* hybrid dysgenesis. For all backcross experiments we used F1 males from TE+ and *D. lummei* crosses. We used these specific F1 males because this genotype was used previously to infer the genetic basis of hybrid dysgenesis within *D. virilis* (Lovoskaya et al. 1990; see below); however interspecific backcross genotypes generated using these males are also the most informative for identifying any additional genic incompatibilities between the TE+ and *D. lummei* parents. To evaluate interspecific isolation and hybrid dysgenesis in parallel, these F1 males were backcrossed individually to either *D. lummei* or TE- females. We evaluated two main classes of models with fitness data from the resulting BC hybrids. Both classes broadly describe Dobzhansky-Muller incompatibilities, because they involve negative epistatic interactions between loci in the two parental strains; however, they differ in whether these interactions occur between genes or between TEs and TE suppressors. We expected that hybrid dysgenesis within *D. virilis* would correspond to the TE model of incompatibility, based on previous work (Lovoskaya et al. 1990; Vieira et al. 1998). Nonetheless, we recapitulated and analyzed data from the dysgenic crosses in parallel to our interspecific crosses, to confirm those prior results.

### Comparing observed patterns to theoretical expectations

Here we describe the unique predictions that are made by different incompatibility models, focusing on two major classes (Fig. 2): models that assume interactions between genes from each parental genome (Genic models) and models that assume an asymmetric copy number of TEs between the parental genomes (TE models). We further summarize the expectations for these models and the data that do and do not support these expectations in Supplemental Table 2.

**Figure 2.**
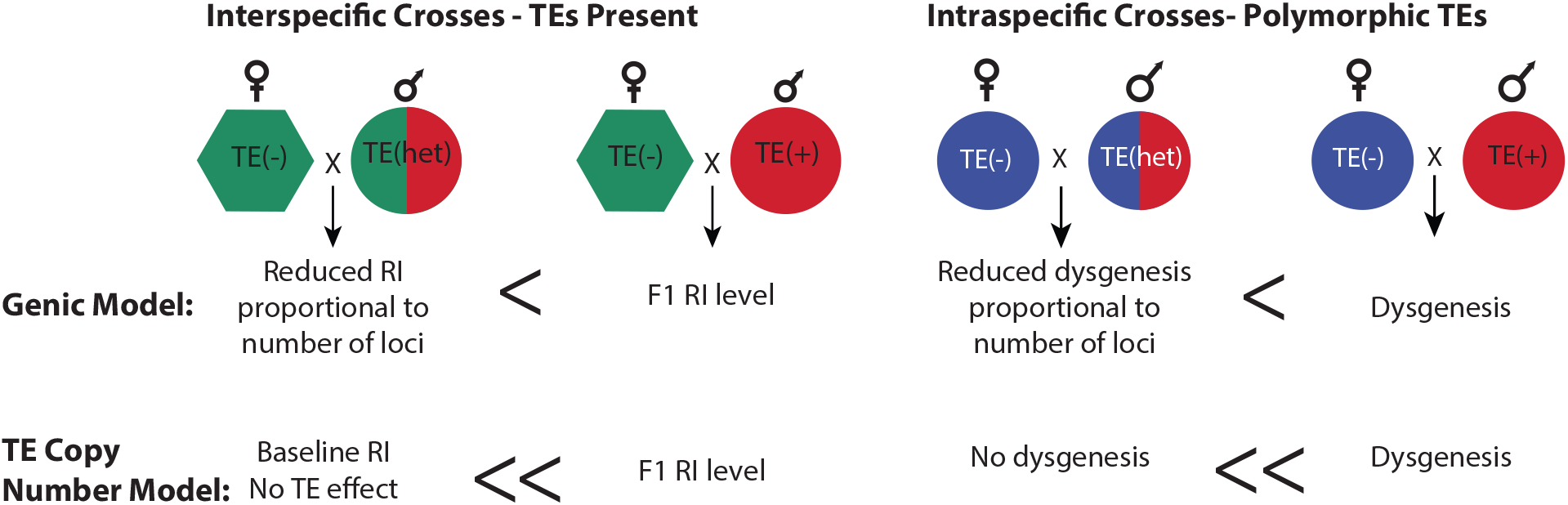
A prediction exclusive to the TE model is that F1 males have reduced TE copy number so their BC offspring will have rescued fertility or viability phenotypes that do not differ from the baseline level of intrastrain crosses. This level of rescue can be distinguished from genic models of hybrid incompatibility.

#### Genic models of hybrid incompatibilities

The first set of models we compared to our empirical observations assume that alternative alleles at incompatible loci have diverged in each lineage and have a dominant negative interaction in F1 and in backcross progeny that carry these alleles (Coyne and Orr 2004). Note that other models of hybrid dysfunction, such as hybrid breakdown, assume negative effects are recessive and only observed in F2 and backcrosses (Coyne and Orr 2004), but here we are interested in incompatibilities that are dominant negative in the F1 individuals. Under our genic model, the proportion of progeny that exhibit the incompatibility phenotype in the backcross will be less than the number/proportion observed in F1 (Fig. 2). That is, backcrossing rescues a proportion of the wild type (viable/fertile) phenotype. The extent of this rescue is determined by the number of interacting loci contributing to the incompatibility, whether these loci are located on autosomes or sex-chromosomes, and the number of independent genetic interactions. For example, when two unlinked autosomal loci cause F1 hybrid incompatibility, we expect two discrete classes of backcross (BC1) phenotypes—half the progeny should carry the incompatible genotype and half should not—and therefore a higher average viability/fertility phenotype. Predictions for the expected level of rescue for several different genic scenarios are described in the Supplemental Methods. This provides an expectation for us to compare observed levels of incompatibility between F1 and BC1 crosses (see Statistical Analysis below).

#### TE models of hybrid incompatibility

To contrast with genic models of hybrid incompatibility we assessed the fit of our observations to a model that is based on TE copy number. Under this model we expect postzygotic reproductive isolation in only one direction of a cross between parental species, specifically when the male parent carries dysgenic TEs but the female parent lacks these (and therefore lacks the production of suppressing piRNAs; Lovozkaya et al 1990; Blumenstiel and Hartl 2005). We also predict a substantial or complete rescue of hybrid fitness in the backcross (Fig. 2), as a result of insufficient dosage of dysgenic causing TEs in these later generation hybrids (Lovoskaya et al. 1990; Vieira et al. 1998). In particular, F1s have half the copy number of dysgenesis causing TEs, so when these F1s are used as the male parent in a backcross to TE- females, TE copy number in their BC offspring can fall below a minimum number necessary to elicit dysgenesis. This dosage/copy number effect for the *D. virilis* system was first suggested by Vieira et al. (Vieira *et al*. 1998) and has also been supported in the *P*-element system in *D. melanogaster* and *D. simulans* (Serrato-Capuchina *et al*. 2020b). Moreover, previous observations indicate that the progeny from crosses between TE- females and males heterozygous for dysgenic factors (i.e., F1 males from TE+ × *D. lummei* crosses) are not dysgenic (Lozovskaya et al. 1990); we interpret this as indicating that the copy number of dysgenic factors (TEs) in the F1 male parent is insufficiently high to cause a dysgenic phenotype in their BC1 offspring. Therefore, while both the TE and genic models predict rescue of wild type progeny in backcrosses, the expected level of rescue from the TE copy number model is significantly higher than predicted from the genic hybrid incompatibility models (Supplemental Methods).

### Phenotypes measured to infer dysgenesis and post-zygotic reproductive isolation

The phenotypes that we measured tracked the developmental time course of hybrid individuals. We first broadly estimated prezygotic isolation to capture potential mating differences or post-copulatory pre-fertilization defects in our crosses; this was measured by comparing the proportion of fertilized and unfertilized embryos from large cage experiments. Our measures of postzygotic isolation included inviability (hatchability, progeny production, and sex ratio) and sterility (testes dysgenesis) in hybrid offspring. The testes dysgenesis phenotype in F1s is characteristic of the intraspecific dysgenic system, but for interspecies incompatibilities we wanted to be agnostic to the type of incompatibility that we might observe, including earlier stages of F1 inviability. These phenotypes were measured in both F1 and backcross experiments and could assess both germline and somatic effects of TEs. Since hatchability and progeny production is the result of both prezygotic success and offspring viability, we could leverage our prezygotic measures to better interpret these measures of inviability in our hybrids.

#### Prezygotic isolation and early stage inviability

From preliminary crosses we knew that the chosen strains all mated and premating isolation was not complete, so we chose to use large crosses in cages to estimate prezygotic isolation and early stage inviability (i.e. hatchability). We set up cages consisting of 25 males and 25 females for each genotype combination (3 replicate populations cages were set up per cross type). Males and females were left in the cages for one week and were given grape agar plates supplemented with yeast culture daily. After the acclimation period, a fresh grape agar plate was given to each cage for embryo collection. We allowed females in the cage to oviposit on this plate for 24 hours and then prepared embryos for staining. Embryos were collected, fixed, and stained with DAPI following basic protocols (Rothwell and Sullivan 2000; Rothwell and Sullivan 2007; Ferree and Barbash 2009). Briefly, embryos were collected in a nitex basket using 0.1% Triton X-100 Embryo Wash (Rothwell and Sullivan 2007). Embryos were dechorionated using a 2.5 % sodium hypochlorite solution (50% solution of household bleach) for 3 minutes and rinsed well with water. Embryos were fixed with 37% formaldehyde and n-Heptane and then vitelline membrane was removed by serially shaking and washing in methanol. To stain embryos, they were first rehydrated by rinsing with PBS and then blocked by incubating with PBT blocking buffer (PBTB), followed by rinsing with PBS before staining with PBTB that contains 1X DAPI for 5 minutes. Embryos were visualized using a fluorescent scope immediately after staining to determine if development had progressed. Embryos were considered to have begun development if we could see nuclei patterns typical of preblastoderm, blastoderm, and gastrula stages. Since we did not score fertilization by co-staining and looking for presence of sperm or whether the second phase of meiosis (formation of polar bodies) had been initiated, we may have missed very early stage embryo lethality and classified these eggs as unfertilized. Thus our estimates of embryo lethality are conservative (they potentially underestimate embryo lethality). All females were dissected after this last embryo collection to confirm they had mated by looking for the presence of sperm stored in the reproductive tract.

To determine if there were differences in viability in early stage embryos we estimated hatchability. After allowing females to oviposit on a fresh grape agar plate for 24 hours, we removed the plate and counted all the eggs. Seventy-two hours later (Day 4) all eggs and larvae on the same plate were counted. We calculated hatchability as the proportion of eggs that failed to hatch/total number of eggs. This was confirmed by counting larvae on the plates. Each hatchability plate was paired with a plate used for the test of fertilization (above).

#### Hybrid progeny production, sex ratio, and sterility

The remaining phenotypes were estimated with progeny generated from individual pairings, rather than population cages. After each male by female pairing was set up, parent individuals were kept for seven days. On day 4 they were transferred to a new fresh vial. On day 7 the parents were transferred to a vial that had been previously dyed with blue food coloring to assist counting of eggs. All vials were cleared and parental individuals discarded at day 8. To ensure that we compared progeny of the same age among different crosses, we did not use any progeny from the first three days of the cross because both intra-strain and inter-strain crosses involving *D. lummei* females would often produce few or no progeny within this time. To be used in further analyses, an individual cross must have produced at least 10 progeny over the 4 day period (days 4-7).

To estimate the total number of progeny produced and their sex ratio, we counted the number of male and female progeny from days 4-7. We determined the viability of individuals for each cross by counting the number of eggs in the day 7 vial as this was the day when females were laying sufficient numbers of eggs to accurately determine hatching rates. We then allowed these eggs to hatch and develop and counted the number of pupae and adults. The number of progeny produced and egg to adult viability gave the same results; since the total number of progeny produced was collected from more replicates and could be summarized by normal distributions we only present those data.

To estimate male gonad atrophy we collected all flies that eclosed from days 4-6 of the cross and saved males into new vials that we supplemented with extra yeast paste, with <10 males per vial. We aged these flies on yeast for five days before scoring gonad atrophy on a dissecting scope by looking for the presence of atrophied testes (Blumenstiel and Hartl 2005). In crosses between Strain 9 (our TE-) and Strain 160 (an inducing strain) it has previously been noted that both, one, or zero testes can be atrophied (Blumenstiel and Hartl 2005). Therefore our estimate for analysis was the proportion of testes atrophied for the total number of testes scored. We also examined female ovary atrophy for a small sample from each interspecies cross but did not observe any atrophy, so we focus exclusively on male gonad atrophy.

### Statistical analysis

All analyses were performed in R v3.3.2.

#### Comparison of embryo development and hatchability measurements

To quantify prezygotic isolation and early embryo lethality (postzygotic isolation), we compared our estimates of embryo development and hatchability using contingency tables. We used the proportion of eggs that had not begun development past the pre-blastoderm stage as our measure of prezygotic isolation. Since there were few eggs for a given embryo collection replicate, we pooled across all three replicates. We then compared the proportion of eggs hatching to the proportion of eggs fertilized using a Chi-square test of independence. We could also compare differences in the proportion of eggs fertilized between crosses that shared a common female genotype, to infer prezygotic (fertilization) isolation.

#### F1 reproductive isolation

For traits measured in the crosses used to generate F1s we performed three separate analyses, based on the identity of the female used in the cross. Since most of these traits are potentially influenced by female fertility, all comparisons were made using the same female parent genotype. To determine if there were differences in trait values based on the male parent in the cross we used linear and generalized linear models. All models had the form

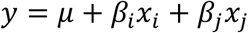

*μ* represents the trait value for the intra-strain cross. *β_i_* and *β_j_* represent the effect sizes for males that were from different strains/species than the female in the specific analysis. For example, when we analyzed data where TE- was the female parent, *β_i_* could represent the deviation caused by having the TE+ strain as the father and *β_j_* could represent the deviation caused by having *D. lummei* as the father. For a given phenotype there was significant reproductive isolation when *β_i_* or *β_j_* was significantly less than zero, indicating lower trait value than the intra-strain cross. In the case of sex-ratio, any deviation of the correlation coefficient from zero could be interpreted as sex specific effects. The dysgenesis and total progeny produced variables were analyzed with simple linear regressions. Since the sex ratio data are more accurately represented by binomial sampling, we analyzed these data using binomial regression (a generalized linear model).

To examine whether there was asymmetry in reproductive isolation for the *D. lummei* and TE+ strain cross, we converted these phenotypes into relative values based on the intra-strain cross. For example, TE+ female × *D. lummei* male measurements can be converted to relative values by finding the difference from the baseline (TE+ intra-strain cross) average value. These relative measurements were compared in pairwise t-tests.

#### Backcross reproductive isolation

The results for the backcross experiment were analyzed in the same way as the F1 crosses. We compared phenotypes for crosses based on the female parent. Specifically, we compared the phenotype values for the control *D. lummei* intra-strain cross with the crosses involving *D. lummei* females mated to TE+ or F1 male genotypes (*D. lummei* comparisons). Similarly, we compared the phenotype values for the control TE- (*D. virilis*) intra-strain cross with the crosses between TE- females and TE+ or F1 male genotypes (*D. virilis* comparisons). The goal of this analysis was to determine if our observed data more closely matched a genic model or TE model of incompatibility. The details of these models are laid out in the Supplemental Methods. To illustrate, we describe one example of their contrasting predictions here. Specifically, for a two locus genic model of incompatibility with an autosome-autosome interaction, when F1 males are used as backcross parents an intermediate number of progeny (regardless of sex) should exhibit the hybrid incompatibility, compared to the parental and F1 crosses. The effect size in this backcross should be ½ of the effect size observed in the original parental cross, regardless of the specific penetrance of the epistatic interaction. For example, for 30% penetrance—that is, 30% of F1 progeny are dysgenic—15% of the backcross progeny are expected to be dysgenic. This contrasts with the TE model where we would expect to see no dysgenesis (i.e. complete rescue) if F1 males as parents lack the number of TEs required to elicit dysgenesis.

For the gonad atrophy data, we analyzed only the data from the F1 male that carried the X chromosome from TE+ since the other genotype was already determined to not be significantly different from zero (see Results). For the data on numbers of progeny produced, we pooled all backcrosses for this analyses since they produced the same number of progeny on average (see Results).

### Data availability

All data and R code used is available through dryad (https://doi.org/10.5061/dryad.zkh1893cj) and Zenodo (https://doi.org/10.5281/zenodo.6567648).

## Results

### Estimates of copy number and divergence among TEs that induce dysgenesis

With our copy number estimates from genomic data, we identified two elements, *Polyphemus* and *Slicemaster*, but not *Penelope*, that had higher average copy number in the inducing *D. virilis* strains—Genome Strain (TE+) and Strain 160—compared to the noninducing *D. virilis* Strain 9 (TE-) and two *D. lummei* strains. Interestingly, we also found copies of *Penelope* in the *D. lummei* genome despite previous reports of the absence of this element (Zelentsova *et al*. 1999). We confirmed that our approach for estimating copy number was consistent with previous approaches that analyzed differences between Strain 160 and Strain 9 (Erwin *et al*. 2015; Funikov *et al*. 2018). Specifically, we found that *Penelope, Polyphemus, Paris, Helena, Skippy*, and *Slicemaster* were enriched in Strain 160 compared to Strain 9 (Fig 3A). Using hierarchical clustering across the *D. virilis* strains to determine which element had a copy number distribution most similar to *Penelope*, we determined that *Polyphermus* and *Penelope* formed a cluster that did not include the other candidate TEs (Fig. 3B). This indicates that *Polyphermus* is the element whose distribution most closely matches that of *Penelope*. The correlation in copy number between *Penelope* and *Polyphemus* across strains was *r*=0.68 (Fig. 3C, *P*=0.0073). When one outlier (Dvir87) was removed the correlation increased to *r*=0.866 (*P*=0.0001). Either correlation indicates a strong relationship between copy number of these two elements and suggests that *Polyphermus* could potentially explain previous copy number dependent dysgenesis in this system.

**Figure 3.**
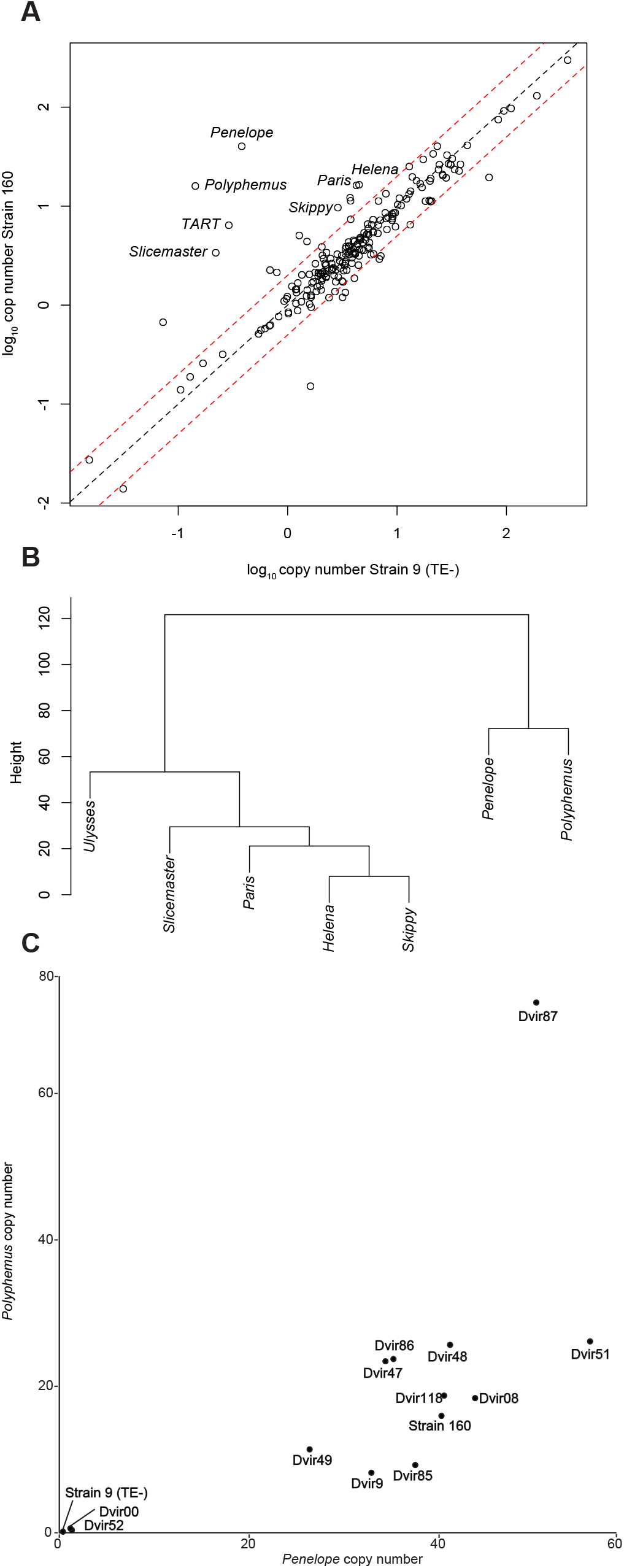
Identification of dysgenic causing transposable elements (TEs) in *D. virilis*. A) Candidate TEs have higher copy number in the inducing Strain 160 compared to the noninducing Strain 9 (TE-). Red dashed line represents 2-fold increase in copy number. B) Clustering based on copy number of candidate TEs across strains of *D. virilis*. C) Positive relationship between the copy number of *Penelope* and *Polyphemus* across *D. virilis* strains.

We used the allele frequencies of single nucleotide polymorphisms (SNPs) in reads mapped to a consensus sequence for each TE to assess the likelihood that *Polyphemus* was active in inducing strains but not in non-inducing strains. We expected active elements to have homogenous insertions with very few SNPs (Erwin et al. 2015). The pattern for *Polyphemus* was consistent with our expectation for the inducing *D. virilis* strains and non-inducing *D. virilis* strain, and for *D. lummei* although there was variation in the estimates between the *D. lummei* samples (Supplemental Fig. 1). While one *D. lummei* sample was nearly identical to the noninducing Strain 9 (TE-), the other strain had a higher copy number and average major allele frequency, however *Polyphemus* was still lower in both *D. lummei* samples than in the inducing strains (Supplemental Fig. 1). We also assessed evidence for activity of *Slicemaster* in inducing strains. In contrast to *Polyphemus*, the major allele frequency data for *Slicemaster* was consistent with a recent broad invasion into the *D. virilis* clade because all strains, with the exception of one *D. lummei* strain, had high average major allele frequency. As a result, for *Slicemaster* there was no consistent pattern differentiating inducing *D. virilis* and non-inducing *D. virilis*, making it an unlikely candidate contributing to dysgenesis.

Combined, our observations from within *D. virilis* and comparison between *D. virilis* and *D. lummei* suggest that copy number differences in *Polyphemus* might play important roles in hybrid dysgenesis. This is consistent with analyses that suggest *Polyphemus* induced DNA breaks in dysgenic progeny (Hemmer *et al*. 2019).

### No postzygotic isolation occurs in interspecies crosses when TEs are absent

We did not observe inviability or gonad atrophy in males or females in either reciprocal cross between TE- and *D. lummei* (Fig.4; Fig. 5.). We did observe an 18% reduction in progeny production in the TE- female × *D. lummei* male cross compared to the intra-strain cross; the former produced, on average, 22 fewer adult progeny than the TE- intra-strain control cross (*μ*=123.318; *P*<0.0001; *β*=-22.509; *P*=0.0341). However, two comparisons indicate that this reduction in progeny production results from prezygotic isolation (specifically reduced fertilization) in this particular direction of the cross, rather than reduced F1 embryo viability. First, the proportion of fertilized embryos did not differ from the proportion of embryos hatching within this cross, indicating no evidence for post-fertilization embryo lethality (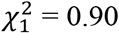, *P*=0.34). Second, the estimated proportion of fertilized embryos was significantly less for the TE- × *D. lummei* cross compared to the TE- intra-strain control cross (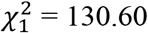, *P*<0.001; Table 2), consistent with a prezygotic fertilization barrier.

**Figure 4.**
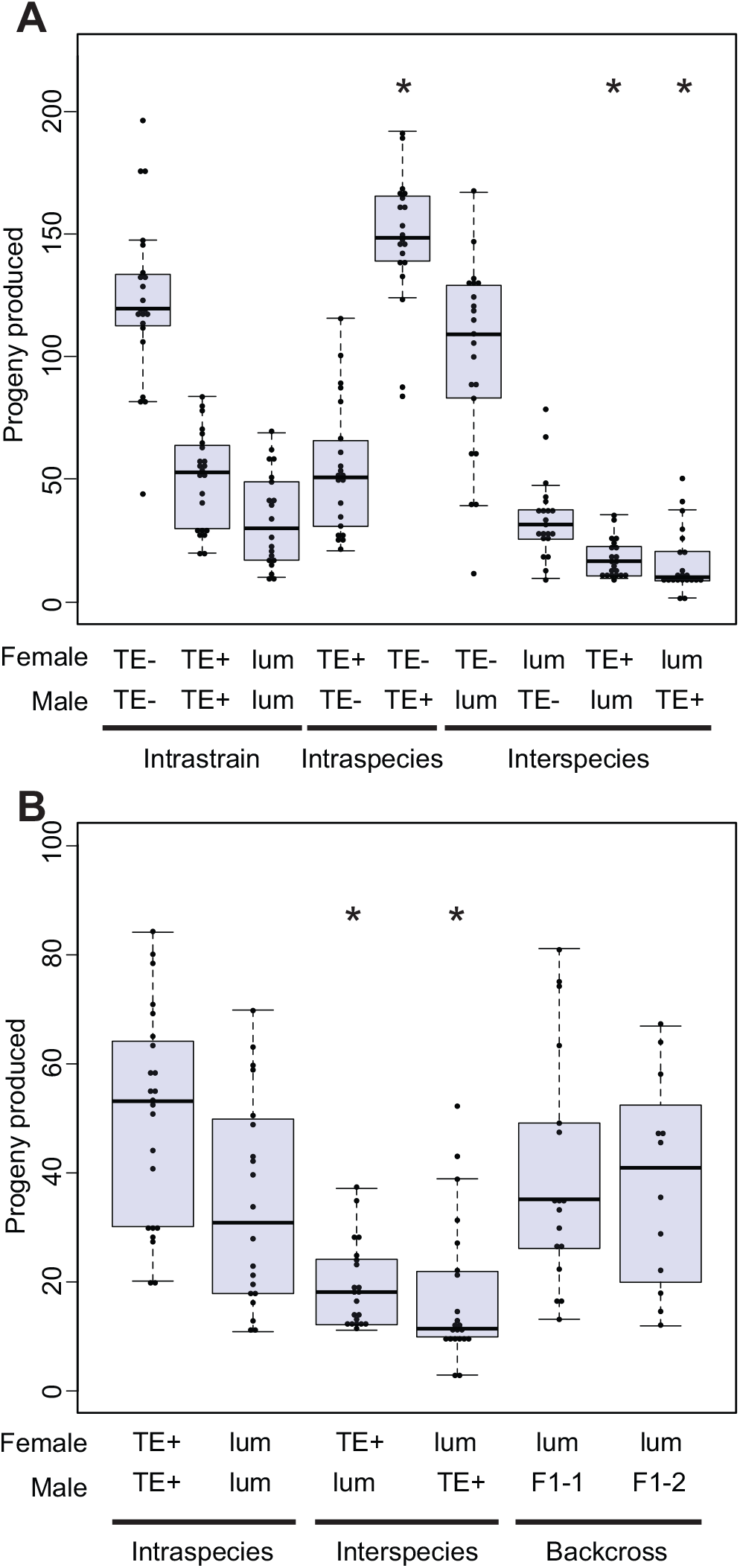
Interspecies crosses with the TE+ strain produce fewer progeny in both directions of the cross combination. A) Intrastrain control crosses, intraspecific crosses, and interspecific crosses. B). Backcross progeny compared to the *D. lummei* intrastrain cross, showing that the F1 males do not induce progeny lethality and progeny production is rescued. F1-1 are males from the *D. lummei* × TE+ cross and F1-2 males are from the reciprocal cross. Each individual dot overlayed on the boxplot is the value for a single replicate. * indicates a regression coefficient that is significantly different from the intrastrain cross of the maternal line using a linear regression model (*P*<0.05).

**Figure 5.**
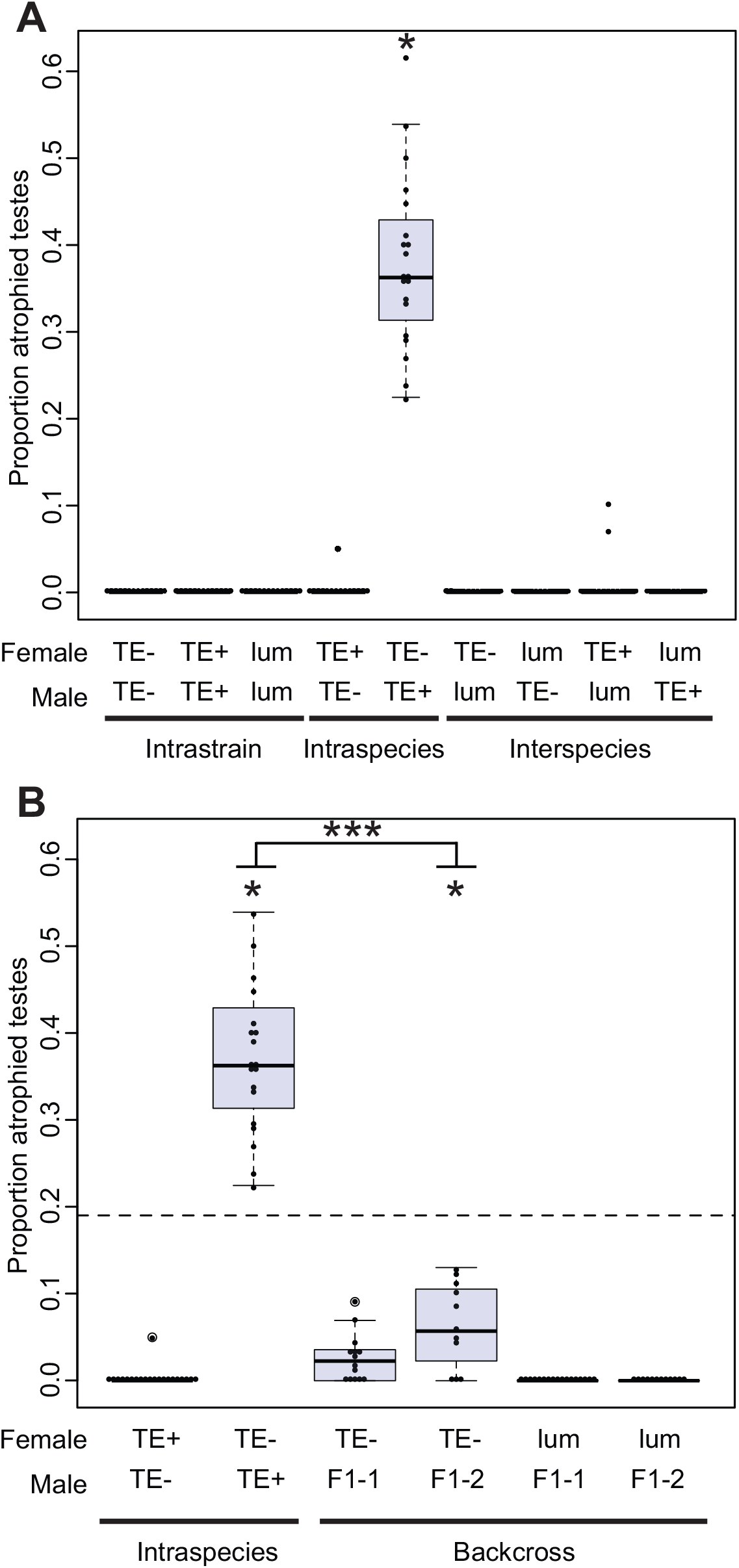
Gonad atrophy characteristic of the classic dysgenic phenotype. A) Intrastrain control crosses, intraspecific crosses, and interspecific crosses demonstrating the lack of dysgenesis except in the intraspecific cross. B) Dysgenesis in backcross progeny compared to TE- × TE+ cross showing that the F1 males do not induce dysgenesis to the same extent as the pure TE+ males. F1-1 are males from the *D. lummei* × TE+ cross and F1-2 males are from the reciprocal cross. Each individual dot overlayed on the boxplot is the value for a single replicate. * indicates a regression coefficient that is significantly different from zero (*P*<0.05) in a linear regression model. *** indicates a significant difference between the progeny means for TE+ and F1 males (*P*<0.001) using a t-test. These regression coefficients also have non-overlapping confidence intervals in a linear model. The dashed line represents the expected effect size based on the two-locus incompatibility models.

The reciprocal cross, *D. lummei* female × TE-, had no evidence of prezygotic isolation. We actually observed a higher proportion of hatched embryos compared to our estimate for proportion fertilized. This observation likely reflects differences in attrition between the techniques used to estimate each stage; in particular, fertilization rate relies on handling embryos with multiple washing and fixing steps which can lead to loss of embryos in the final sample (Rothwell and Sullivan 2007b).

### Strong F1 postzygotic isolation occurs in interspecies crosses when TEs are present

In contrast to the TE- interspecific crosses, we observed significantly fewer progeny in both directions of cross between the TE+ strain and *D. lummei* (Fig. 4), compared to intra-strain (control) crosses. In the *D. lummei* female × TE+ male cross, where we would expect to see the effects of TEs, 17 fewer progeny were produced compared to the *D. lummei* intra-strain cross (μ =34.50; *P*<0.001; *β*=-16.95; *P*=0.002). This was also significantly lower than the number of progeny produced in the *D. lummei* female × TE- male interspecific cross, where TEs are absent (*D. lummei* female × TE+ male 95% Confidence Interval= −27.86, −6.04; *D. lummei* female × TE- male 95% Confidence Interval = −10.25, 11.82). We found that the reciprocal TE+ female × *D. lummei* male cross also produced fewer progeny compared to the intra-strain (TE+) control cross (mu=50.52; P<0.0001; *β*=-31.37; *P*<0.0001); in this case 19 fewer progeny. In both reciprocal crosses involving *D. lummei* and TE+, the proportion of embryos hatched was significantly lower than the proportion of embryos fertilized, indicating that the reduced number of progeny was due to early embryo lethality in both cases (*D. lummei*×TE+, 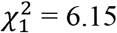, *P*=0.01; TE+×*D. lummei*, 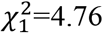, *P*=0.02; Table 2).

In only one cross direction did we observe sex specific inviability. In the TE+ female × *D. lummei* male cross, males had increased inviability, resulting in a significant excess of females and a sex ratio that was 63% female (*β*=0.1171; *P*<0.0001; Fig. 6). Finally, while the cross between TE+ females and *D. lummei* males produced rare male gonad atrophy, we conclude that gonadal dysgenesis occurs mainly in the intraspecific TE- female × TE+ male cross (Fig. 5).

**Figure 6.**
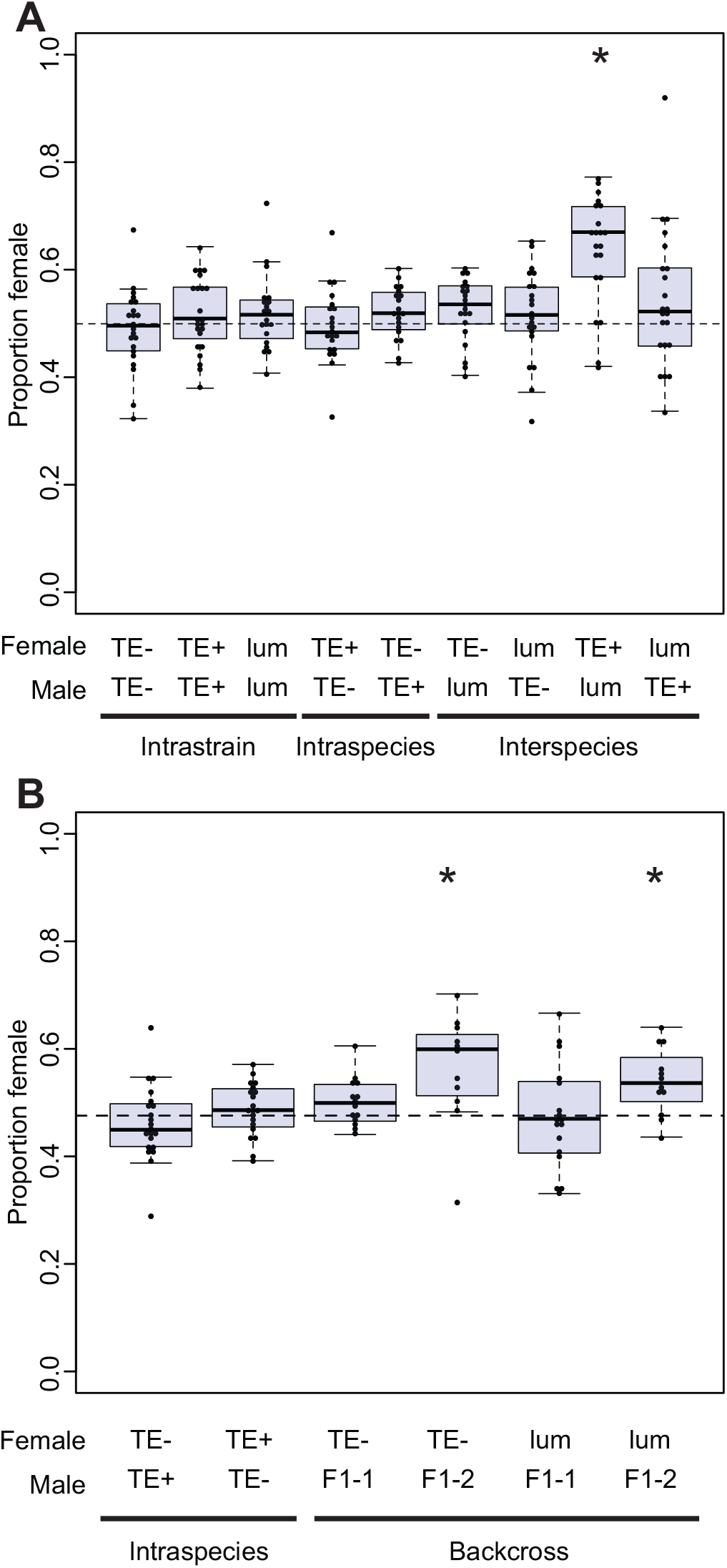
The sex ratio of each cross used to examine deviations from a 50:50 sex ratio caused by sex specific lethality. A) Intrastrain control crosses, intraspecific crosses, and interspecific crosses. B). The sex ratio of backcross progeny showing sex biased progeny ratios occur when *D. lummei* Y is in combination with some complement of TE+ autosomes. F1-1 are males from the *D. lummei* × TE+ cross and F1-2 males are from the reciprocal cross. Each individual dot overlayed on the boxplot is the value for a single replicate.* indicates significant deviation from 50:50 sex ratio (*P*<0.05) using a binomial regression model.

### Backcrosses indicate that both TEs and genic interactions contribute to F1 hybrid incompatibility phenotypes

Together our data for F1s suggested that there are possibly two mechanisms contributing to hybrid incompatibilities in these crosses. In particular, the TE model predicts asymmetric reproductive isolation, specifically reduced F1 fitness in the *D. lummei* female × TE+ male cross. Therefore, our observation of F1 inviability in both crossing directions suggests an additional mechanism contributing to postzygotic isolation in the TE+ male *× D. lummei* female cross. Alternatively, the observed F1 hybrid inviability in both directions could be due to a genic autosome-autosome mechanism acting in both cross directions, without a contribution from TEs. We were able to differentiate between these (and more complex) models using our series of backcrosses.

In these experiments, we examined progeny production and sex ratio in backcrosses between *D. lummei* females and F1 males that were created in both reciprocal directions between *D. lummei* and the TE+ strain. Both models make predictions for the expected rescue in progeny number/viability and for sex specific lethality in the BC when F1 males from either initial crossing direction are used as the male parent. Under the TE copy number model, we expect complete rescue of progeny number and no sex specific lethality effects. The genic models predict rescue of varying magnitude depending on the number of interacting loci; however, for up to 4 interacting loci, the expected rescue is always less for the genic incompatibility models compared to the TE copy number model. For example, under an autosome-autosome or X-autosome genic model with two interacting loci, we expect an intermediate number of BC progeny compared to the (dysgenic) *D. lummei* female × TE+ male cross and the (control/non-dysgenic) intrastrain *D. lummei* × *D. lummei* control cross, unlike complete rescue under the TE copy number model. In both backcrosses the observed number of progeny in our experiments was not significantly different than the *D. lummei* × *D. lummei* control cross, and represented a complete rescue of viability, consistent with the TE copy number model (Fig. 4). To quantitatively test the predictions from the genic models, we used a series of *t-tests* where the expected number of progeny produced was determined by the assumed number of interacting loci. These tests indicate that the rescue we observe does not fit any models of genic incompatibility involving up to 4 interacting loci (Table 3; Supplemental Materials). Combined, these tests consistently indicate that the magnitude of BC rescue we observed fits the TE copy number model better than a genic model of incompatibility (Table 3; Fig 4).

**Table 3.** Progeny production is rescued when F1 males are crossed to *D. lummei* females. For the backcrosses we tested whether progeny rescue was greater than the expectation for two locus genic incompatibility models (26 progeny produced). F1-1 are males from the *D. lummei* × TE+ cross and F1-2 males are from the reciprocal cross. For the progeny produced we report the number of replicate crosses (n) and the standard error (se). *indicates a significant difference in progeny production from the intrastrain control.

| Female | Male | Mean Progeny Produced (n, se) | t statistic (df) | P-value |
| --- | --- | --- | --- | --- |
| *D. lummei* | *D. lummei* | 34.50 (20, 4.28) | NA | NA |
| *D. lummei* | TE+ | 17.54* (22, 2.78) | NA | NA |
| *D. lummei* | F1-1 | 39.76 (17, 5.26) | 2.61 (16) | 0.009 |
| *D. lummei* | F1-2 | 38.41 (12, 5.56) | 2.22 (11) | 0.023 |

Finally, analyzing the sex ratio in these backcrosses also allowed us to identify an autosome-Y incompatibility that can explain, in part, the inviability observed in F1s from the TE+ female × *D. lummei* male cross, where we do not expect to see an effect of TEs. In particular we observed a significant excess of females (*β* =0.25; p=0.037) in the backcross involving *D. lummei* females and F1 males that carried the X chromosome from TE+ (Fig. 6). Neither our TE copy number model nor the specific X-autosome incompatibility model we evaluated predict sex-specific effects past the F1 generation (see methods). The only two other crosses where we observed a skewed sex ratio were also female-biased and involved F1 males that carried the X chromosome from TE+ and the Y chromosome from *D. lummei*, crossed with *D. lummei* (*β*=-0.252, *z*=-2.08, *P*=0.037) or TE- females (*β*=-0.273, *z*=-3.491, *P*<0.001). In all three of these cases, males had different X chromosome genotypes but consistently had the *D. lummei* Y chromosome (Fig. 6) From this, we conclude that this additional incompatibility is caused by an autosome-Y interaction.

### The classic intraspecific dysgenesis system conforms to the TE copy number model

We expected that the classic dysgenic cross (TE- female × TE+ male) within *D. virilis* would also be consistent with a TE copy number model, based on previous work (Lozovskaya et al. 1990 Vieira et al. 1998). Nonetheless, we recapitulated the originally reported crosses and interspecific backcrosses to confirm this expectation and to compare our inferences in parallel with interspecific hybrid incompatibility.

As expected, in the offspring of TE- females crossed to TE+ males we observed 38% testes atrophy (Fig. 5), a frequency that was significantly greater than the TE- intra-strain cross (*β*=0.3807; *P*<0.0001); in the reciprocal non-dysgenic cross (TE+ female × TE- male) we observed rare gonad atrophy events at a frequency not significantly different from zero. We did not observe inviability in either reciprocal cross. The non-dysgenic cross produced significantly more offspring than the TE- female × TE- male intra-strain comparison (*β*=26.732; *P*=0.0136); this increased productivity may reflect some outbreeding vigor between these long-established lab lines. There was no deviation in the proportion of eggs hatching compared to the proportion fertilized in either cross, indicating there was no embryo inviability (TE-×TE+, 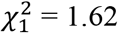, *P*=0.20; TE+×TE-, 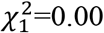, *P*=1.00; Table 2).

As with our interspecific crosses, to differentiate whether the observed patterns of dysgenesis were more consistent with a TE model or a genic incompatibility model we used a directed backcross experiment using the same F1 male genotypes used in previous analyses (Lovavskaya et al. 1990) and in our interspecific experiments. For the TE model we expected to observe rescue of gonad atrophy in a backcross using F1 males (as outlined above, and Supplemental Methods), compared to our baseline level of gonad atrophy observed in F1 males of the dysgenic (TE- × TE+) cross—i.e., 38%. We found that this rescue varied with the specific X chromosome inherited in the BC, but was nonetheless consistent with strong rescue of fertility predicted from the TE copy number model. When F1 males carried the X chromosome from *D. lummei* the level of dysgenesis was not significantly different than zero (*β*=0.0264; *P*=0.1366). When F1 males carried the X chromosome from TE+ the level of dysgenesis was ~6%, also significantly less than the dysgenic cross (*β*=0.0638; *P*=0.0011; CI= 0.0261, 0.1016) and less than 19% atrophy that would be expected from a genic model.

## Discussion

In this study, we used a powerful set of crosses to test whether the presence of TEs influences the magnitude of reproductive isolation. Using the *D. virilis* intraspecific dysgenic system we predicted that the *D. virilis* strain carrying inducing TEs should show greater postzygotic reproductive isolation when crossed with *D. lummei* lacking these TEs, compared to the cross involving the *D. virilis* strain that lacks inducing TEs. We indeed observed this pattern of elevated postzygotic isolation specifically in the TE+ interspecific cross, along with other more nuanced differences between our focal crosses. Taken together, our genetic data from both F1 and backcross experiments indicate that a model where TEs contribute to elevated reproductive isolation in the interspecific cross fits better than alternative models of genic incompatibilities. In addition, with genomic data we also assessed which candidate TE might be responsible for observed dysgenesis patterns among inducing *D. virilis* strains compared to non-inducing *D. virilis* and *D. lummei*, and confirmed previous reports that *Polyphemus* is a strong candidate for causing dysgenesis (Funikov *et al*. 2018; Hemmer *et al*. 2019).

Although a role for selfish genetic elements in the expression of postzygotic reproductive isolation is consistent with the Dobzhansky-Muller model of hybrid incompatibility (Johnson 2010; Castillo and Moyle 2012; Crespi and Nosil 2013), the difficulty of explicitly disentangling TE effects from the effects of other hybrid incompatibilities is underappreciated. For example, one way TEs are thought to contribute to reproductive isolation is through transcriptional misregulation that causes sterility or inviability (Martienssen 2010; Michalak 2010; Dion-Cote *et al*. 2014). However, this is not a unique feature of TE-based hybrid incompatibility; divergence in trans and cis-regulatory elements are common and can also cause misregulation and hybrid incompatibilities, via ordinary genic effects (reviewed in Mack and Nachman 2017). To disentangle TE-specific from these other effects it would also be necessary to differentiate whether TE misregulation is simply symptomatic of more general misregulation in hybrids or if TE divergence itself drives global misregulation. Instead of focusing on analyzing patterns of misregulation, here we took the approach of isolating phenotypic effects due to TEs by using strains with defined differences in TE copy number, and potential activity, making explicit predictions about which hybrid classes should exhibit increased reproductive isolation, and evaluating these using classical genetic crosses.

The strength of this approach relies on our ability to make and test predictions that are exclusive to a TE incompatibility model, compared to alternative genic models, in parallel for two crosses. We observed that postzygotic incompatibility occurred only in crosses involving the TE+ carrying *D. virilis* strain, and not in the cross between TE- *D. virilis* and *D. lummei*, clearly implicating TEs in this postzygotic isolation. We were able to support this inference by using backcrosses, and comparing these results to parallel crosses examining intraspecies dysgenesis. A key prediction, specific to the TE incompatibility model, arises from the observation that F1 males fail to produce dysgenic sons when crossed to TE- females (Lovoskaya et al. 1990); the underlying mechanism we propose, based on previous studies (Vieira *et al*. 1998; Srivastav and Kelleher 2017; Serrato-Capuchina *et al*. 2020b), is that F1s have sufficiently low copy number of TEs such that dysgenesis does not occur when F1 males are themselves used as fathers. This pattern allowed us to directly compare the TE incompatibility model with genic incompatibility models because the former consistently predicts greater rescue of incompatibility in the BC generation, irrespective of the specific incompatibility phenotype observed. We found that crosses between F1 males and TE- females and F1 males and *D. lummei* females produced largely concordant results. Both incompatibility phenotypes—atrophied testes in crosses between TE- *D. virilis* females and F1 males, and viability in crosses between *D. lummei* females and F1 males—were completely rescued. Both intra- and inter-specific results are consistent with TE- mediated incompatibility. Results from these backcrosses also enabled us to determine the basis of additional, non-TE, interactions contributing to male-specific inviability observed exclusively in the TE+ × *D. lummei* cross. We infer that this sex-specific effect results from an autosome-Y incompatibility. Interestingly, autosome-Y incompatibilities have been previously reported for hybrid sterility in the *D. virilis* clade (Lamnissou *et al*. 1996; Heikkinen and Lumme 1998; Sweigart 2010); a role for TEs in this phenotype has not been examined although specific models of TE regulation (e.g. the location of a protective piRNA cluster on the TE+ Y-chromosome) could potentially explain such autosome-Y effects in hybrids. Regardless, together our observations support the operation of two distinct mechanisms of hybrid inviability between *D. lummei* and *D. virilis*, one of which directly implicates TEs in the expression of this postzygotic isolating barrier.

A second inference from our findings is that TEs can cause different hybrid incompatibility phenotypes—that is, sterility or inviability—depending on context. In particular, our observations suggest that both sterility and inviability (in intra- and inter-specific TE+ crosses, respectively) result from misregulation of the same TEs at different developmental stages. Because TE misregulation can cause DNA damage leading to cell death and tissue atrophy it has the potential to produce multiple different incompatibility phenotypes (Tasnim and Kelleher 2018; Phadnis N. *et al*. 2015). Moreover, although TEs have typically been thought to restrict their damage to the germline, this is not always the case (Borque *et al*. 2018), including in *D. virilis* system where germline and somatic expression have been demonstrated for *Penelope* (Blumentstiel and Hartl 2005). Direct evidence that TEs that can cause inviability phenotypes when active in somatic tissue comes from a *P-element* mutant engineered to have expression in somatic cells (Engels *et al*. 1987); crosses involving this somatically-expressed *P-element* mutant line showed inviability that was dependent on *P*-element copy number. These observations indicate it is mechanistically plausible that dysgenic elements can be responsible for both inviability and sterility phenotypes, as suggested by our findings, although the specific connection between damage and these phenotypes needs further exploration. The involvement of TEs in both germline and somatic effects in hybrids could also provide a direct mechanistic connection (via TE activity) between intraspecies dysgenesis and interspecies reproductive isolation.

Finally, our observations also identify a mechanism that could contribute to variation among intraspecific strains in the strength of their isolation from other species. Evidence for polymorphic incompatibilities is typically inferred from the observation of among-population variation in isolation phenotypes, when crossed with a second species. The specific alleles underlying this variation are usually unknown (Reed and Markow 2004; Kozlowska *et al*. 2012), but these polymorphic incompatibility loci are typically assumed to be based on genic Dobzhansky-Muller incompatibilities (Cutter 2012). In this study, we show that the presence/absence of TEs segregating among populations and species can also contribute to intraspecific variation in the strength of isolation between species. Moreover, the rapid pace with which TE copy number and identity can change within a lineage means that such differences could rapidly accumulate between species, especially species pairs in which there is known to be dynamic turnover in the identity of their active TEs.

Overall, here we provide evidence for a role of TEs in increased reproductive isolation between lineages that differ in the presence of specific active TE families, by leveraging known TE biology to differentiate TE vs non-TE effects in a set of targeted crosses. The generality of these results could be assessed by focusing on other *a priori* cases where species are known to vary in the number or identity of TEs. For instance, increased reproductive isolation caused by TEs was recently reported in crosses between *D. simulans*, that are polymorphic for *P-elements*, and a sister species, *D. sechellia*, that completely lack *P-elements* (Serrato-Capuchina et al. 2020a). Similar studies that examine lineages at a range of stages of TE differentiation could address the contribution of TEs to the accumulation of hybrid incompatibilities over evolutionary time, especially in comparison to ‘ordinary’ genic incompatibilities. At present, our and other recent (Serrato-Capuchina et al. 2020a), findings suggest that the rapid lineage-specific evolution of TEs could potentially explain the frequent observation of polymorphic incompatibilities in species that experience dynamic changes in selfish genetic elements, and could therefore represent one of the earliest arising mechanisms of postzygotic reproductive isolation in these groups.

