## Supplemental Methods and Information for "Hybrid incompatibility between *D. virilis* and *D. lumei* is stronger in the presence of transposable elements"

**Supplemental Information**

**Supplemental Methods**

**Genic models of hybrid incompatibilities**

We generated both qualitative and quantitative expectations to determine whether the data are more consistent with a genic model of incompatibility or a TE copy number model. Qualitative differences between the TE model and genic models are described in the main text. Here we provide more detail on the complete set of models that we compared. Our genic models vary in complexity but still demonstrate that even under conditions where you would expect to see the greatest phenotypic rescue in backcrosses—when incorporating many interactions for genic incompatibilities—the TE model is still a better fit for the observed data.

For a genic incompatibly involving two unlinked autosomal loci we expect to see discrete classes of backcross (BC1) genotypes that exhibit hybrid incompatibilities because half of the progeny should carry the incompatible genotype and half should not (Supplemental Fig. 2). If the hybrid incompatibility is not fully penetrant then, on average, any specific replicate BC1 cross would have less half the number of progeny exhibiting the hybrid incompatibility as were observed in F1s from the original parental cross between species. For example, if in the F1 cross the proportion of inviable progeny was 30%, then the proportion of inviable progeny in the BC1 would be 15%. This assumes the penetrance is consistent between these genotypes, and the segregation of alleles results in half the number/proportion of affected progeny.

We expanded this general model to include X-autosome interactions among two or more loci. In particular, we considered an X-autosome model where there is a negative interaction between the X chromosome of the TE- or *D. lummei* parent and an autosome of the TE+ parent (Supplemental Fig. 3). We restricted ourselves to this specific X-autosome interaction because this generates dysgenesis/incompatibility in both sexes for the canonical intraspecific dysgenic cross (TE- × TE+), but no male gonad dysgenesis/incompatibility in the reciprocal cross, consistent with previous observations of the intraspecific dysgenesis phenotype (Lozovskaya et al. 1990). For models in which higher order (greater than 2 loci) interactions are responsible for the incompatibility observed in the F1s, we would expect to observe a smaller proportion of affected individuals in the backcross compared to the F1 cross (Supplemental Table 2); as shown in the table, the specific expected degree of hybrid incompatibility in the backcross depends on the total number of loci required for hybrid incompatibility and the number of loci carried by each parental strain.

More complex models of incompatibilities can be generated, but a realistic upper limit on the number of interactions must take into account how many could segregate in the backcross, given the lack of recombination in F1 *Drosophila* males used as backcross parents. For example if there are four loci that contribute to two independent incompatibilities in the F1, then on average the effect seen in the backcross progeny will be 1/2 of the F1 effect (Supplemental Fig. 4), which is not different than the predictions from our less complex models. Similarly, loci that are physically linked can also contribute to reductions in the effect of DMIs. In the same example of four loci and two incompatibilities, if two of the parental alleles are physically linked with no recombination, the backcross effect is also 1/2 of the F1 effect (Supplemental Fig 5). This would phenotypically be indistinguishable from the two locus model. More complicated models—for example, where incompatibilities accumulate at very different rates within alternative lineages—can be produced, but many are not plausible within the biological reality of the system. For example, the difference between F1s between *D. virilis* TE (+) and *D. lummei*—where we see clear hybrid incompatibility— and F1s between *D. virilis* TE (-) and *D. lummei* —where we see no evidence of incompatibilities (either sterility or viability)—could be explained by all of the incompatible alleles arising in the TE(+) lineage only. However, given that the TE(+) and TE(-) are lab strains that have likely only been separate since right before the most recent invasion of *Penelope,* the rapid accumulation of genic incompatibilities exclusively in this single strain is very unlikely.

**Supplemental Results**

Two-locus genic incompatibility models might be an oversimplification when considering hybrid incompatibilities (main results Table 3), so we also considered models with higher order interactions (greater than two independent/freely-recombining loci). For a three-locus autosomal interaction the greatest expected rescue would be 1/4 of the original effect. With our data, therefore, we would expect 9.5% dysgenesis in crosses between backcross males and TE- females. For crosses between backcross males and *D. lummei* females the phenotype is different (the number of progeny produced) and applying the 1/4 effect would result in the expectation of ~30.26 progeny produced. Using a one sample t-test we rejected the hypothesis that our data can be explained by a three-locus incompatibility for gonadal dysgenesis in the backross male crossed to TE- female (H0: $\mu<9.5$; $t= -627.2$, $df = 10$, $P< 0.0001$) and for number progeny produced (H0: $\mu>30.26$; $t=2.36$, $df=28$, $P=0.0126$) in the backcross male crossed to *D. lummei* female. For a four-locus autosomal interaction, the largest rescue would be 1/8 the original effect corresponding to 4.75% dysgenesis in crosses with TE- females and 32.38 progeny produced in crosses with *D. lummei* females (Supplemental Table 2). A one-sample t-test also rejected the hypothesis for the four-locus incompatibility for gonadal dysgenesis (H0: $\mu<4.75$; $t= -311.48$, $df = 10$, $P<0.0001$) and number progeny produced (H0: $\mu >32.38$; $t=1.802$, $df=28$, $P=0.041$). We did not explore interactions beyond four loci because (as explained above) *D.*

*virilis* only has four major autosomes and, since we used F1 males that lack recombination, each chromosome is transmitted as a single locus in our crosses.

**Supplemental Table 1.** SRA accession information for the genomic sequence data used in make estimates of transposable element copy number.

| SRA accession | Strain Name | Species Stock Center # | Additional info |
|---|---|---|---|
| SRR1200631 | Strain 160 | NA. Dr Justin Blumenstiel | inducing strain |
| SRR1200817 | Strain 9 (TE-) | NA Dr Justin Blumenstiel | non-inducing strain |
| SRR5278980 | Genome Strain (TE+) | 15010-1051.87 Dr Andy Clark | inducing strain |
| SRR9261305 | Dvir85 | 15010-1051.85 | Sendai, Japan. |
| SRR9261306 | Dvir52 | 15010-1051.52 | U.S.S.R. (1976). |
| SRR9261307 | Dvir51 | 15010-1051.51 | Santiago, Chile. neutral strain (Vieira et al 1998) |
| SRR9261308 | Dvir49 | 15010-1051.49 | Chaco, Argentina (1950). |
| SRR9261310 | Dvir9 | 15010-1051.09 | Sendai, Japan |
| SRR9261311 | Dvir86 | 15010-1051.86 | Puebla, Mexico (1947). neutral strain (Vieira et al 1998) |
| SRR9261316 | Dvir87 | 15010-1051.87 | Genome Strain |
| SRR9261322 | Dvir47 | 15010-1051.47 | Hangchow, China. neutral strain (Vieira et al 1998) |
| SRR9261323 | Dvir48 | 15010-1051.48 | Puebla, Mexico (1947). neutral strain (Vieira et al 1998) |
| SRR9261328 | Dvir08 | 15010-1051.08 | Truckee, California. |
| SRR9261329 | Dvir118 | 15010-1051.118 | Gikongoro, Rwanda (2009) |
| SRR9261331 | Dvir00 | 15010-1051.00 | Pasadena CA |
| SRR5278981 | Dnova4 | 15010-1031.04 | Moab, Utah (1949) |
| SRR5278982 | Dlum8 | 15010-1011.08 | Hokkaido, Japan. |
| SRR5278983 | DamML975 | NA. Dr Yasir Ahmed-Braimah | NA |
| SRR9426109 | Dlum9 | 15010-1011.09 | Moscow |
| SRR9426111 | Dnova00 | 15010-1031.00 | Grand Junction, Colorado |
| SRR9426112 | Dam00 | 15010-1041.00 | Anderson, Indiana. |

**Supplemental Table 2**: Summary of predicted patterns of reproductive isolation from interspecific crosses, based on either Genic or TE Models of hybrid incompatibility, and observed experimental results used to assess these predictions. Note that predictions and observed data are for interspecific crosses only.

| | Mechanism of Interspecific RI | Difference between reciprocal crosses (F1 generation) | Sex-specific F1 hybrid incompatibility | Fitness of BC hybrids from F1 fathers | Sex-specific BC hybrid incompatibility |
|---|---|---|---|---|---|
| Predictions from: | | | | | |
| Genic Models* | | | | | |
| • Autosome-autosome interaction | Interaction between at least one autosomal locus from each parental genome | None predicted (Supplemental Figure 2) | None predicted (Supplemental Figure 2) | Partial rescue: Intermediate** between F1 and intrastrain fitness (Supplemental Figure 2, 5, 6) | None predicted (Supplemental Figure 2) |
| • X-Autosome interaction (where autosome is from TE+ parent) | Interaction between at least one X-linked locus and one autosomal locus from alternative parental genomes | Yes—stronger isolation predicted from cross direction in which maternal lineage carries the X-linked incompatible allele, compared to reciprocal cross direction (Supplemental Figure 3) | Yes—isolation phenotype predicted in females from cross in which paternal lineage carries the X-linked incompatible allele (Supplemental Figure 3) | Partial rescue: Intermediate between F1 and intrastrain fitness (Supplemental Figure 3) | None predicted for this specific X-A model (Supplemental Figure 3) |
| TE Model | Asymmetric copy number of TEs (and suppressing piRNAs) between parental genomes | Yes—isolation phenotype predicted in TE- female x TE+ male cross, but not in reciprocal cross direction | None predicted | Full rescue: no difference from intrastrain fitness | None predicted |
| Observations from experimental data: | | | | | |
| (Progeny production (days 4-7)) | | No—Progeny production (due to embryo lethality) is reduced in both cross directions (Figure 3A, Table 2) | | Full rescue: no difference in progeny production compared to intrastrain fitness (Figure 3B, Table 3) | |

| | | | | | |
|---|---|---|---|---|---|
| Offspring gonadal atrophy | | No substantial atrophy detected (Figure 4A) | | No substantial atrophy detected (Figure 4B) | |
| Sex-ratio bias (female bias due to male inviability) | | Yes—Reduced male viability in TE+ female x D. lummei cross only (Figure 5A) | Yes—Reduced male viability in TE+ female x D. lummei cross only (Figure 5A) | | Yes—Reduced male viability in BCs from F1 males carrying D. lummei Y-c'some only (Figure 5B) |

*These genic models address interactions that can cause dominant negative interactions in F1s, as this is the primary phenotype we seek to explain in our crosses. ** The magnitude of partial rescue depends on the number of loci contributing to the incompatibility (see Supplementary methods, and Supplemental Table 2).

**Supplemental Table 3.** Summary of possible Dobzhansky-Muller hybrid incompatibilities involving different number of loci and different numbers of loci per parental strain. If the effect size in F1 individuals is "h" the proportion of progeny with incompatible genotype is reduced in the backcross (a fraction of h). These interactions assume all dominant autosomal interactions. This scenario represents the F1 backcrossed to Parent 2. Each loci is bi-allelic with alleles being lower case or upper case.

| **Two locus interaction** | | | |
|---|---|---|---|
| Parent 1 | Parent 2 | Incompatible genotype | Proportion BC progeny with HI |
| AAbb | aaBB | A-B- | 1/2 h |
| **Three locus interaction** | | | |
| Parent 1 | Parent 2 | Incompatible genotype | Proportion BC progeny with HI |
| AAbbCC | aaBBcc | A-B-C- | 1/4 h |
| AAbbcc | aaBBCC | A-B-C- | 1/2 h |
| **Four locus interaction** | | | |
| Parent 1 | Parent 2 | Incompatible genotype | Proportion BC progeny with HI |
| AAbbCCDD | aaBBccdd | A-B-C-D | 1/8h |
| AAbbCCDD | aaBBCCdd | A-B-C-D | 1/4 h |
| AAbbccdd | aaBBCCDD | A-B-C-D | 1/2 h |

**Supplemental Figure 1.** Copy number and average major allele frequency for candidate dysgenic TEs demonstrating activity across the *virilis* clade. A) The pattern for *Penelope* is used as reference since it has previously been used as an example to demonstrate strains with average major allele frequency close to 1 are active. B) The pattern for *Polyphemus* is similar to *Penelope* and demonstrates that this element is likely not active outside of *D. virilis* inducing strains. C) The pattern for *Slicemaster* contrasts with the other elements suggesting recent invasion in the clade. For all plots Strain 160 and GS (TE+) are inducing strains. Strain 9 is a non-inducing strain. Dvir86, Dvir47, Dvir48, and Dvir51 are "neutral" strains (Vieira et al 1998). Dnov represents *D. novamexicana*, Dam represents *D. americana*, and Dlum represents *D. lummei*

**Supplemental Figure 2.** Graphical representation of an autosome-autosome incompatibility model between the genomes of the TE(-) and TE(+) strains.

**Supplemental Figure 3.** Graphical representation of an X-autosome incompatibility model, highlighting that even though sex specific effects are predicted in the F1 non-dysgenic cross, both backcrosses should produce inviable progeny at ½ the rate of the dysgenic cross.

**Supplemental Figure 4.** Depiction of the reduction in hybrid incompatibilities in the backcross generation compared to the F1 generation when there are two independent incompatibilities contributing to hybrid incompatibility.

**Supplemental Figure 5.** Depiction of the reduction in hybrid incompatibilities in the backcross generation compared to the F1 generation when there are two independent incompatibilities contributing to hybrid incompatibility, but there is linkage between loci.

### Supplemental Figure 1.

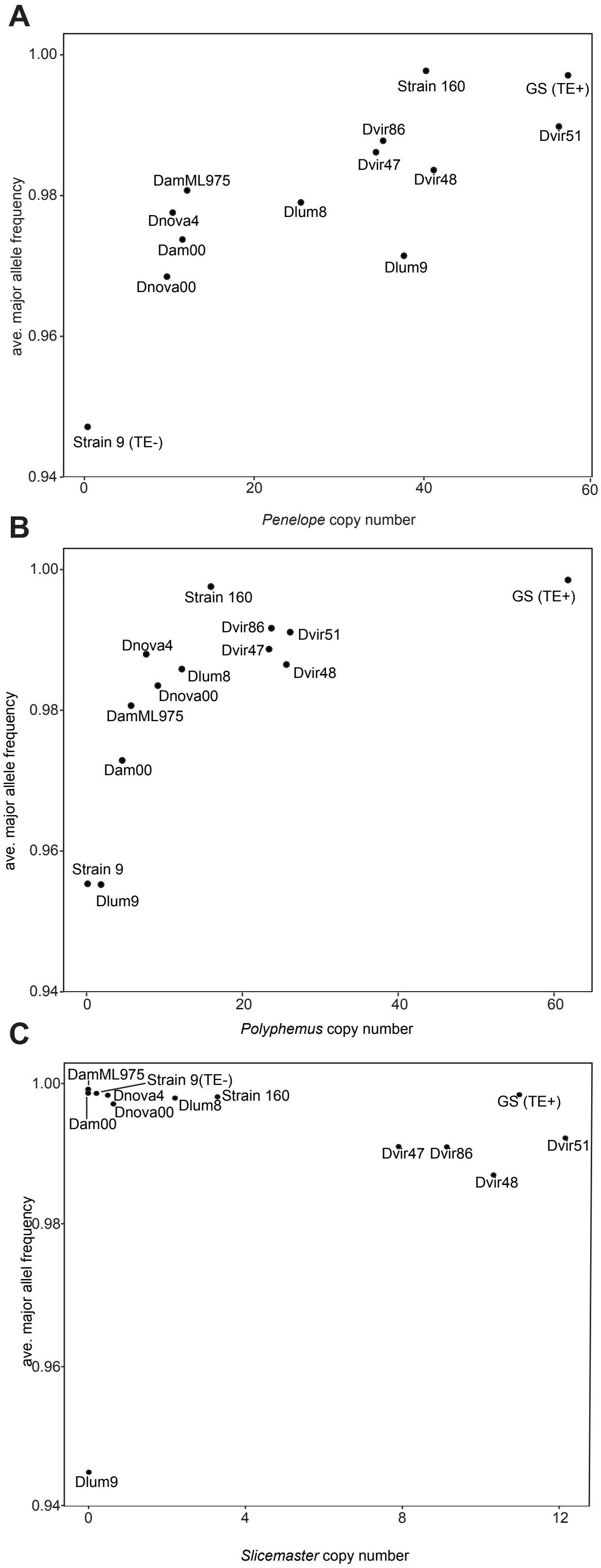

**Supplemental Figure 2**

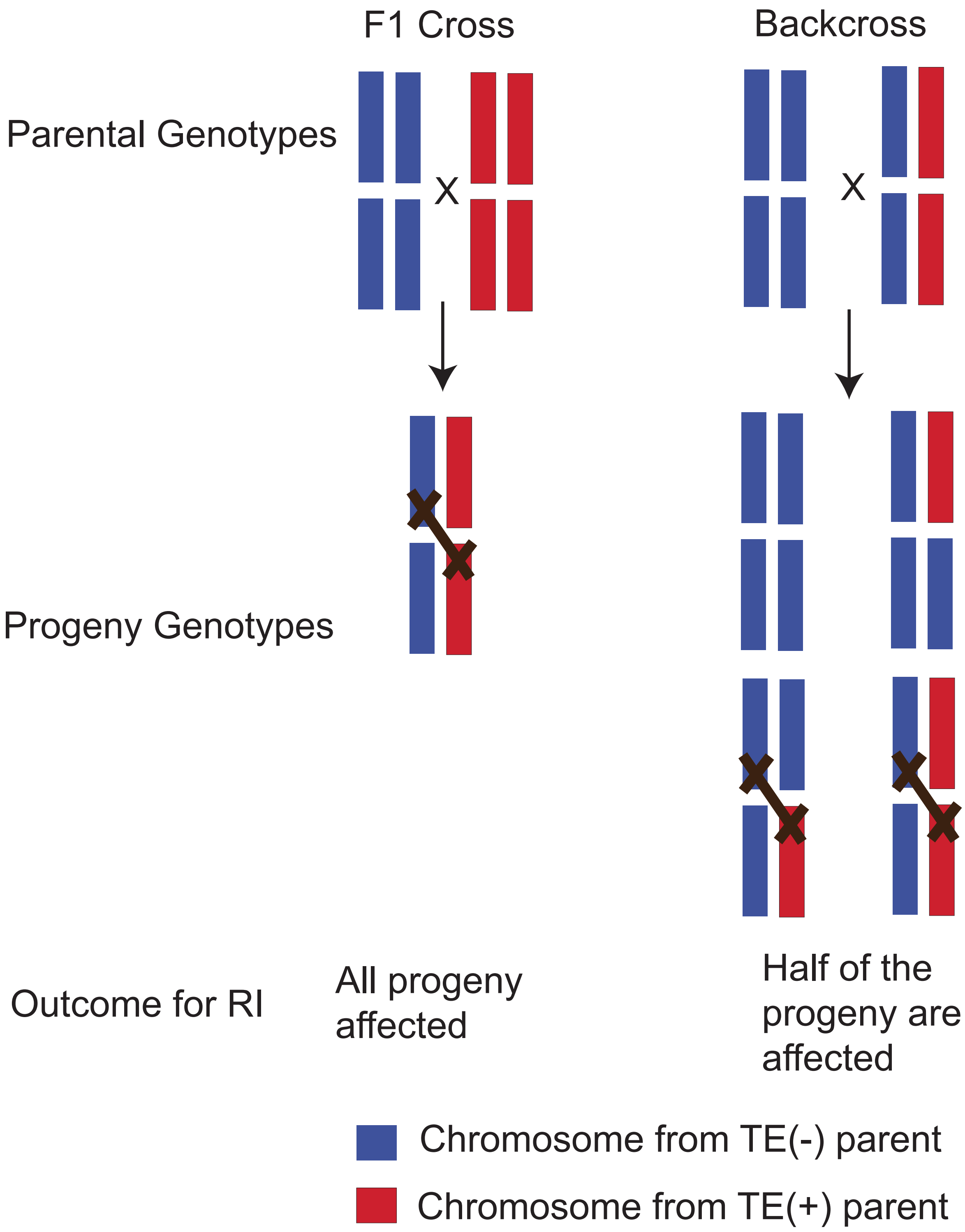

**Supplemental Figure 3**

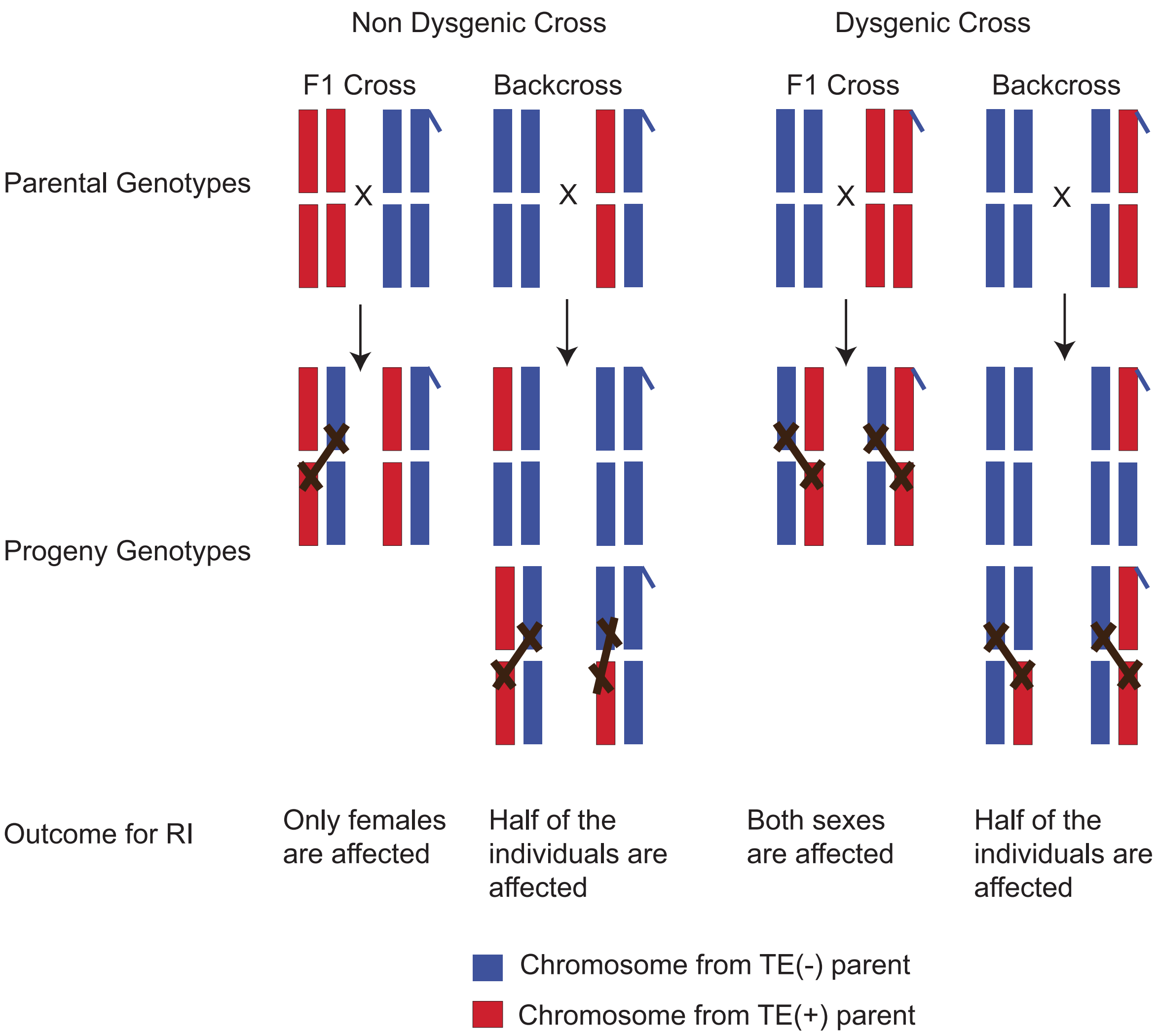

**Supplemental Figure 5**

Cross to generate F1

AAbbCCdd x aaBBccDD

↓

AaBbCcDd

AxB=20%

CxD=20%

Total=40%

Assume independent DMIs between A and B and C and D.

F1 backcrossed to one parent

AaBbCcDd x AAbbCCdd

↓

¼ progeny will have A-B- c-d-=20%

¼ progeny will have a-b- C-D-=20%

¼ progeny will have A-B- C-D-=20%

¼ progeny will have a-b- c-d-=0%

Average effect is 15%

**Supplemental Figure 6**

Cross to generate F1

AACCbbdd x aaccBBDD

↓

AaCcBbDd

AxB=20%

CxD=20%

Total=40%

Assume independent DMIs between A and B and C and D. But the A and C loci are physically linked with no recombination

F1 backcrossed to one parent

AaCcBbDd x AACCbbdd

↓

All progeny will have A-C-

¼ progeny will have A-C-B-D-=40%

¼ progeny will have A-C- B-d-=20%

¼ progeny will have A-C- b-D-=20%

¼ progeny will have A-C- b-d-=0%

Average effect is 20%
